# HIGHLY PHAGOCYTIC LIPID-ASSOCIATED MACROPHAGES (LAMs) ARE INCREASED IN COLONIC LAMINA PROPRIA IN OBESITY

**DOI:** 10.1101/2022.12.20.521239

**Authors:** Angela Castoldi, David E Sanin, Nikki van Teijlingen Bakker, Christiane F Aguiar, Lauar de Brito Monteiro, Nisha Rana, Katarzyna M Grzes, Agnieszka M Kabat, Jonathan Curtis, Alanna M Cameron, George Caputa, Tiago Antônio de Souza, Fabrício O Souto, Joerg M Buescher, Joy Edwards-Hicks, Erika L Pearce, Edward J Pearce, Niels Olsen Saraiva Camara

## Abstract

Little is known about the effects of high fat diet (HFD)-induced obesity on resident colonic lamina propria (LP) macrophages (LPMs) function and metabolism. Here, we report that obesity and diabetes resulted in increased macrophage infiltration in the colon. These macrophages exhibited the residency phenotype CX3CR1^hi^MHCII^hi^, and were CD4^−^TIM4^−^. During HFD, resident colonic LPM exhibited a lipid metabolism gene expression signature that overlapped that used to define lipid associated macrophages (LAMs). Via single cell RNA sequencing, we identified a sub-cluster of macrophages, increased in HDF, that were responsible for the LAM signature. Compared to other macrophages in the colon, these cells were characterized by elevated glycolysis, phagocytosis and efferocytosis signatures. CX3CR1^hi^MHCII^hi^ colonic resident LPMs had fewer lipid droplets (LD) and decreased triacylglycerol (TAG) content compared to equivalent cells in lean mice, and exhibited increased phagocytic capacity, suggesting that HFD induces adaptive responses in LPMs to limit bacterial translocation.

## Introduction

The low-grade inflammatory response in obesity initially results from excessive nutrient accumulation that culminates in altered metabolic homeostasis. Sustained subsequent systemic inflammation involves changes in gut microbiota and increased gut permeability, which are associated with the maintenance of immune cell activation status, and associated comorbidities seen in obese individuals [1-5].

Intestinal inflammation in obese subjects and in mice is subtle compared to that seen in adipose tissue. In obesity, there are broadly more infiltrating immune cells within the lamina propria (LP) [6]. This is consistent with the fact that obesity triggers innate immune responses with increased interleukin (IL)-1β and adaptive immune response with increased T helper 1 (Th1) and Th17 cells in both the small and large intestine [6-11]. Despite the fact that there are changes in the immune cell compartment of the intestine during obesity, it remains unclear whether resident macrophages within this tissue play a role in the inflammatory state. In the majority of tissues, resident macrophages are derived from embryonic precursors that populate the tissue before birth and maintain themselves by self-renewal in the tissue during adulthood, with only minor contributions from blood monocytes. The intestine is an exception to this rule, since the macrophage population (MHCII^hi^CX3CR1^hi^) in this organ requires constant replenishment by blood monocytes throughout adulthood [12]. A small pool of self-maintaining CD4^+^TIM4^+^ macrophages was reported to exist in the adult intestine in mice; these persist independent of replenishment by blood monocytes, but their number declines with age [13, 14]. Another population of CD4^+^TIM4^−^ macrophages with a slow turnover from blood monocytes and a CD4^−^TIM4^−^ population completely dependent on blood monocyte replenishment can also be found in mouse LP [13]. As monocytes enter the LP, they undergo a differentiation process whereby they first express major histocompatibility complex II (MHCII), followed by F4/80, CD64 and CX3CR1 [15, 16]. Intestinal resident macrophages have a role in maintenance of tissue homeostasis, inflammation and inducing resolution after inflammation [16]. During colitis, the terminal differentiation of monocytes into mature resident macrophages (MHCII^hi^CX3CR1^hi^) is disrupted [16] and evidence suggest a causal link between defects in resolution of intestinal inflammation and altered monocyte-macrophage differentiation, with the accumulation of cells that are considered to be immature macrophages causing impaired bacterial clearance and excessive cytokine secretion in patients with inflammatory bowel disease (IBD) [17, 18].

Colonic macrophages have been found to play an important role in the induction of insulin resistance under HFD. CCR2 expression and inflammation were found to be increased in the colon after 12 weeks of HFD feeding, corroborating increased monocyte recruitment to this site, and blocking monocyte, and thereby macrophage recruitment into the colon improved metabolic parameters [19]. However, the role of the other populations of macrophages in the colon during HFD has not been explored. Lipid associated macrophages (LAMs), a subset of macrophages defined by a distinct lipid metabolism-associated signature, have recently been described to play a role in regulating gains in adiposity during obesity [20]. LAMs are increased in adipose tissue during obesity, and recognized to arise from circulating monocytes. They characteristically express Trem2, a sensor of extracellular lipids which is involved in phagocytosis, lipid catabolism and energy metabolism. LAMs were found to benefit systemic metabolism, preventing adipocyte hypertrophy, systemic hypercholesterolemia, inflammation and glucose intolerance. Since the original description, LAMs have been described in several additional disease states, including non-alcoholic steatohepatitis (NASH), where they were found in aggregates in hepatic crown-like structures and were associated with protection against liver fibrosis [21]. In atherosclerosis, LAMs are thought to contribute to the calcification of atherosclerotic lesions [22]. Moreover, LAMs appear to be associated with the pathogenesis of Alzheimer Disease [23]. In breast cancer, LAMs were distributed near adipocytes and characterized by an M2-like activation signature and were noted to be highly phagocytic, and their depletion was associated with protective anti-tumor effects [24]. Despite this growing realization that LAMs are present in a variety of diseases and associated with, depending on context, both positive and negative outcomes, these cells have not been studied in the LP.

In the present study, we report that obesity increases macrophage infiltration in the colon, and that in this setting macrophages acquire the residency CX3CR1^hi^MHCII^hi^ phenotype. Furthermore, we show that the CX3CR1^hi^MHCII^hi^ macrophages that populate the colon in obesity are mainly CD4^−^TIM4^−^. We found that during HFD, CX3CR1^hi^MHCII^hi^ colonic resident LPMs display an increased lipid metabolism signature. Additionally, we found that the CX3CR1^hi^MHCII^hi^ colonic resident LPMs have decreased lipid droplet (LD) accumulation and decreased triacylglycerol (TAG) content, which is associated with increased fatty acid oxidation. CX3CR1^hi^MHCII^hi^ colonic resident LPMs showed increased phagocytic capacity in HFD-fed mice, suggesting that increased lipid metabolism is an adaptive response of these cells to limit bacterial translocation and maintenance of gut homeostasis. Further, through single cell RNA sequencing (scRNAseq), we identified a cluster of macrophages expressing genes consistent with the LAM signature, including genes associated with increased phagocytosis and efferocytosis. Our data indicate that LAMs develop within the lamina propria during HFD.

## Results

### HFD feeding increases colonic resident CX3CR1^hi^ macrophages infiltration

We investigated whether obesity induced by HFD affects colonic LPMs. We used CX3CR1^+/gfp^ mice fed normal chow (Lean) or high fat diet (HFD; 60% fat) for 12 weeks. Compared to lean mice, HFD-fed mice gained a significant amount of fat mass (Supplementary Figure 1a), had increased glucose intolerance, decreased insulin sensitivity (Supplementary Figure 1b), LPS detectable within peripheral blood (Supplementary Figure 1c), and short colons (Supplementary Figure 1d). The total number and frequency of CD64^+^CD11b^+^ colonic LP macrophages were increased because of HFD (Figure 1a-c). The majority of cells in the broad CD64^+^CD11b^+^ population were CX3CR1^hi^MHCII^hi^ LPMs (Figure 1d-f). This suggests that the majority of newly infiltrating macrophages acquire the expression of residency markers.

**Figure 1:**
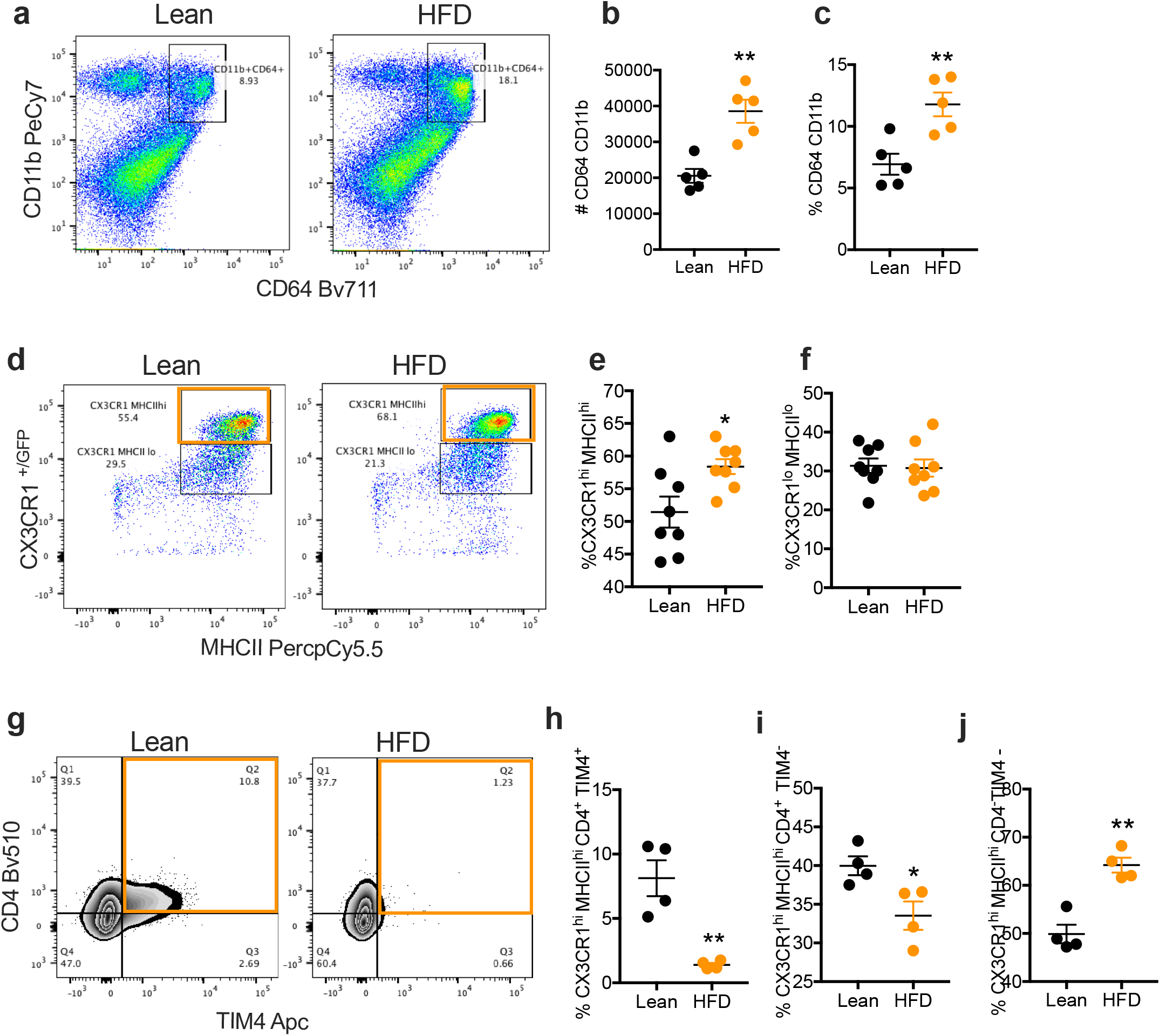
HFD increases macrophage infiltration in the colonic lamina propria. **(a-c)** Analysis by flow cytometry of global macrophages (CD64 CD11b) in the LP of lean and HFD-fed mice. Showing the total number **(b)** and the percentage **(c). (d)** Analysis of MHCII CX3CR1 macrophages showing increased MHCII^hi^ CX3CR1^hi^ **(e)** and no difference in CX3CR1^lo^ MHCII^lo^ **(f)**. Flow cytometry analysis of long lived CD4 TIM4 LPMs gated on CX3CR1^hi^MHCII^hi^ LPMs. Percentage of long lived CD4+ TIM4+ LPMs. **(i)**Percentage of long lived CD4+ TIM4-LPMs. **(j)** Percentage of long lived CD4-TIM4-LPMs.

### Long-lived CD4^+^TIM4^+^ resident LPMs are depleted upon HFD

Recently, a population of long-lived macrophages expressing CD4 and TIM4 was shown to populate the intestinal LP [13]. This population was described to be depleted from the LP in aged mice [13]. We found that HFD feeding resulted in a decrease in the CX3CR1^hi^MHCII^hi^CD4^+^TIM4^+^ population from the colonic LP (Figure 1 g-h). There was also a decrease in CX3CR1^hi^MHCII^hi^CD4^+^TIM4^−^ cells (Figure 1g, i). The resident macrophages that populate the colonic LP on HFD are mainly CX3CR1^hi^MHCII^hi^ CD4^−^TIM4^−^ (Figure 1g, j). Together, our data show that HFD increases CX3CR1^hi^MHCII^hi^ LPMs in the colon, moreover, the long lived CD4^+^TIM4^+^ macrophages are depleted and CD4^−^TIM4^−^ macrophages become the dominant population in mice fed HFD.

### Colonic CX3CR1^hi^ MHCII^hi^ LPMs display a lipid associated phenotype and increased activation state

We sorted CX3CR1^hi^ MHCII^hi^ LPMs from lean and HFD-fed mice and used RNAseq to ask whether HFD leads to changes in gene expression. We found that 167 genes were regulated by > one (1) fold change (Supplemental Figure 2a). Amongst these differentially expressed genes, 40 were up-regulated and 127 genes were down-regulated (Supplemental Figure 2). Further analysis was restricted to the most significantly (p≤0.01) differentially regulated genes (Figure 2a). Genes in this category indicate that HFD promotes changes in the expression of genes that are engaged in inflammation, such as *Tlr2 [25], Vcam1 [26]*, and the negative regulator of IL-1 signaling, *Il1r2 [27]*, and genes related to lipid metabolism, such *Lpl [28], Abcg1 [29], Ppargc1a [30]* and *Aqp9* [31] (Figure 2a-f). The lipid-metabolism associated genes are among genes expressed by LAMs [20], suggesting that HFD induces the LPMs to resemble LAMs.

**Figure 2:**
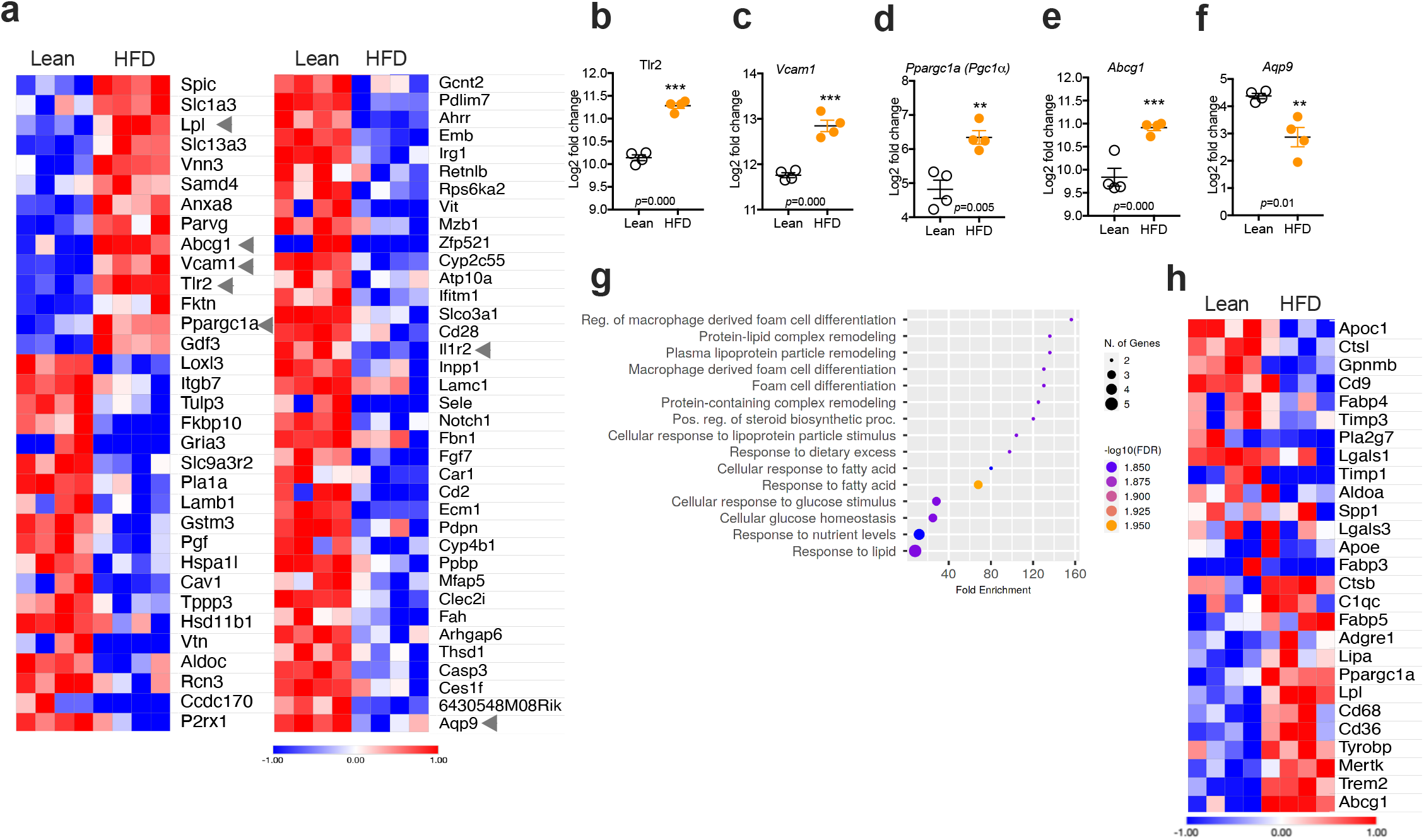
HFD modulates gene expression in the colonic CX3CR1^hi^ MHCII^hi^ LPMs. **(a)** RNA sequencing analysis of the most significantly regulated genes (p<0.01) in CX3CR1^hi^ MHCII^hi^ LPMs from lean and HFD fed mice. Comparison of **(b)** *Tlr2*, **(c)***Vcam1*, **(d)** *Ppargc1a*, **(e)** *Abcg1*, **(f)** *Aqp9* Log2 Fold change between Lean and HFD. **(g)** Fold Enrichment analysis of top15 biological processes related to the up-regulated (p<0.01) genes in CX3CR1^hi^ MHCII^hi^ LPMs from HFD fed mice. **(h)** Heat map of Lipid metabolism related genes in CX3CR1^hi^ MHCII^hi^ LPMs from lean and HFD fed mice. ∗∗p < 0.01; ∗∗∗p < 0.001.

Enrichment analysis based on hypergeometric distribution followed by false discovery rate (FDR) correction for GO terms using the 14 up-regulated genes (p≤0.01), showed enriched biological processes related to lipid metabolism (Figure 2g). Heatmaps of LAM-associated gene [20] expression showed regulation in HFD fed compared to lean mice (Figure 2h).

Moreover, the *RNAseq* data provided support for the flow cytometry-based finding that HDF leads to a decrease in LP CD4^+^TIM4^+^ macrophages (Fig. 1g), revealing decreased *Timd4* expression in LPMs sorted from HFD fed mice while, the expression of *Cd4* and the macrophage markers *Cx3cr1, Fcgr1* and *Lyz2*, were not affected (Supplemental 2b-g).

In parallel, we took a targeted approach using gene expression and flow cytometry to measure the effects of HFD on the expression of known markers of macrophage activation. While expression of *Cd86, Cd80* and *Mrc1* by colonic CX3CR1^hi^MHCII^hi^ LPMs was unaffected (Fig. 3a-c), the surface expression of the proteins that these genes encode, CD80, CD86 and CD206, was increased by HFD-feeding mice (Fig. 3d-j). A similar pattern was observed for CX3CR1^lo^MHCII^lo^ LPMs (Supplementary Figure 3).

**Figure 3:**
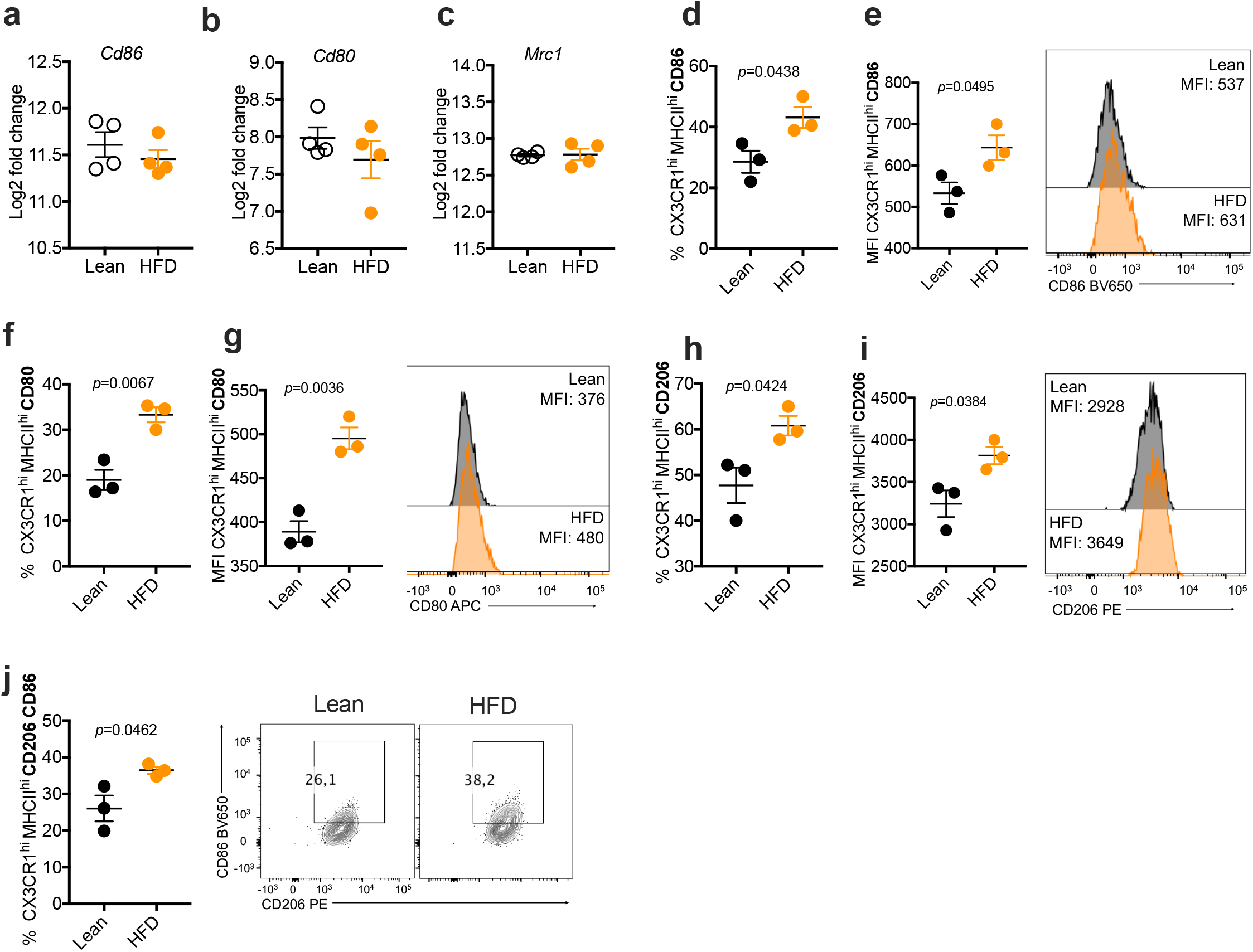
HFD modulates CD86, CD80 and CD206 in CX3CR1^hi^ MHCII^hi^ LPMs. **(a)** Percentage of CX3CR1^hi^ MHCII^hi^ LPMs expressing CD86 and **(b)** MFI of CD86. **(c)**Percentage of CX3CR1^hi^ MHCII^hi^ LPMs expressing CD80 and **(d)** MFI of CD80. **(e)**Percentage of CX3CR1^hi^ MHCII^hi^ LPMs expressing CD206 and **(f)** MFI of CD206. **(g)**Percentage of CX3CR1^hi^ MHCII^hi^ LPMs expressing CD206 CD86. **(h)** Log2 Fold change of gene expression of *Cd86, Cd80* and *Mrc1* by CX3CR1^hi^ MHCII^hi^ LPMs between Lean and HFD. ∗∗p < 0.01; ∗∗∗p < 0.001.

### Single cell profile of colonic LP macrophages after HFD feeding

We used scRNAseq of sorted CD45^+^ colonic LP cells to analyze HFD-induced immune changes in the colon. LP cells were isolated from a total of 6 mice (3 HFD and 3 Lean) and loaded onto the 10X Genomics Chromium platform. After sequencing, aggregation of the samples, quality control, removal of contaminating CD45-cells, and exclusion of cells resembling doublets, a total of 8,207 cells remained (4,613 cells from lean, 3,594 cells from HFD). Seventeen transcriptionally distinct clusters of cells were identified by generating a UMAP from the transcriptome data using principal component analysis (Supplemental Figure 4a). Within these clusters, we identified distinct cell types based on the differentially expressed genes (DEGs) (Supplemental Figure 4a-b). We did not detect different clusters between HFD and Lean conditions (Supplemental Figure 4 b).

**Figure 4:**
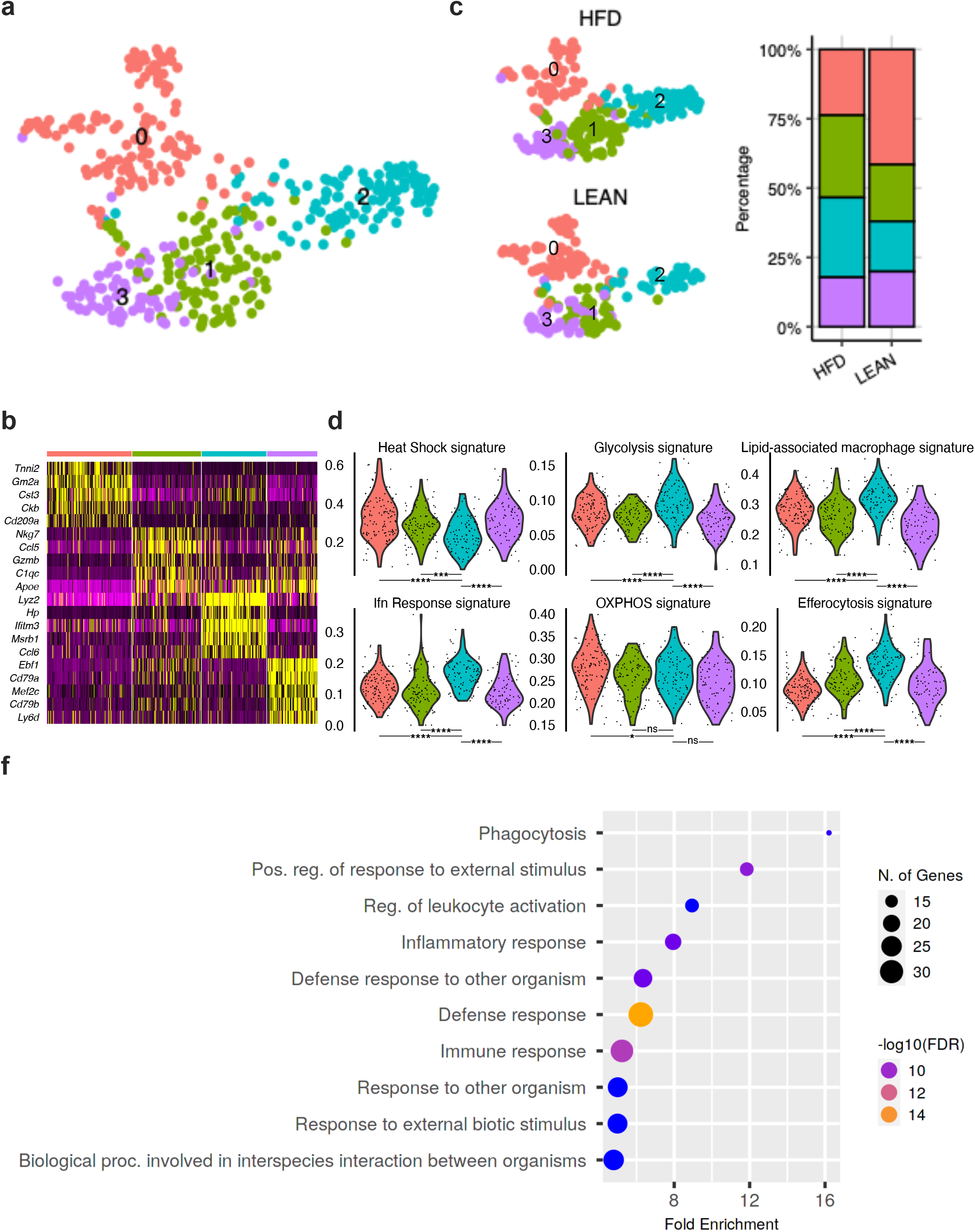
HFD increases the proportion of colonic macrophages with a LAM signature. **(a-b)** Single cell RNA sequencing of CD45+ cells from the LP isolated from 6 mice (3 (12W) HFD and 3 lean) and loaded onto the 10X Genomics Chromium platform. Macrophage Cluster (Cluster 5) was identified according to the expression of *Lyz2, Cx3cr1, Fcgr1, Cd68* and *Adgre1* and re-clustered in a new UMAP divided in 4 clusters. **(c)** Percentage of each cluster in lean and HFD condition. **(d)** Gene expression scores calculated using UCell **(e)** Enrichment analysis of top10 biological processes related to the up-regulated (p<0.01) genes in cluster 2. ∗p < 0.05; ∗∗∗∗p < 0.0001.

Using an unbiased cell classification based on scGate annotation [32] we identified clusters with Lymphoid cell signature, T cell signature, CD4^+^ T cell signature, T regulatory cell signature, CD8^+^ T cell signature, B cell signature, NK cell signature, Myeloid cell signature, and Macrophage signature (Supplemental Figure 4c). The cluster classified by scGate annotation as Macrophages was further shown to express the macrophage-defining genes *Lyz2, Cx3cr1, Fcgr1, Cd68* and *Adgre1*, indicating that cluster 5 are likely macrophages (Supplemental Figure 4d). To examine changes in this population, we re-clustered cells within cluster 5 into a new UMAP, in which four clusters were identifiable (Figure 4a-b). At this resolution, differences in proportion between HFD and lean conditions were evident, with cluster 0 smaller and cluster 2 larger in HFD (Figure 4c). In order to understand the differences between these clusters, we next calculated gene expression scores using the UCell [33] metric to analyze the function of these cells (Figure 4d). Cluster 2, which is expanded in HFD fed mice had a decreased heat shock signature, strong Interferon response and glycolysis signatures and no difference, compared to other clusters, in the overall OXPHOS signature. Corroborating the bulk RNA sequencing data indicating increased lipid metabolism in resident colonic CX3CR1^hi^ MHCII^hi^ LPMs (Figure 2a-h), we found a significantly higher LAM signature (Figure 4 d). Moreover, an increased efferocytosis signature was also detected in cluster 2 (Figure 4d). Based on significantly different gene expression (p<0.01) in cluster 2, we applied enrichment analysis based on hypergeometric distribution followed by false discovery rate (FDR) correction for GO terms (Figure 4f). The 10 top regulated biological processes in cells in cluster 2 included phagocytosis, inflammatory/immune response and defense response (Figure 4f). *Trem2*, a specific LAM marker, was expressed mainly by cluster 2 in the colonic LPMs (Supplemental Figure 4e). This suggests that HFD leads to an increase in in the LP of a population of macrophages that resembles LAMs with an activated phenotype, associated with increased glycolysis, phagocytosis and efferocytosis.

### The neutral lipid content of colonic resident macrophages is reduced during HFD

Our data indicate that HFD induces changes in the expression of genes linked to lipid metabolism in colonic LPMs. To explore this further, we analyzed the lipid content of colonic CX3CR1^hi^ MHCII^hi^ LPMs. Using Bodipy staining, we observed a decrease in neutral lipids/lipid droplets in cells from HFD mice (Figure 5a). Consistent with this, lipidomics indicated that compared to cells from lean conditions, colonic CX3CR1^hi^ MHCII^hi^ LPMs from HFD mice have lower levels of stored triacylglycerols (TGs) (Figure 5b), especially TG-48 and TG-50 (Supplementary Figure 5), although the total lipid content of these cells tended to be higher (Figure 5c and Supplementary Figure 5). We reasoned that the latter may be accounted for by increased expression of *Cd36* (Figure 2h), a major mediator of cellular fatty acid uptake [34]. Further, we reasoned that the observed decrease in TG content in colonic CX3CR1^hi^MHCII^hi^LPMs from HFD mice might be explained by increased fatty acid oxidation (FAO) coupled to TG hydrolysis. Consistent with this, CPT1a expression measured by flow cytometry was increased in the LP macrophages from mice on HFD (Figure 5d).

**Figure 5:**
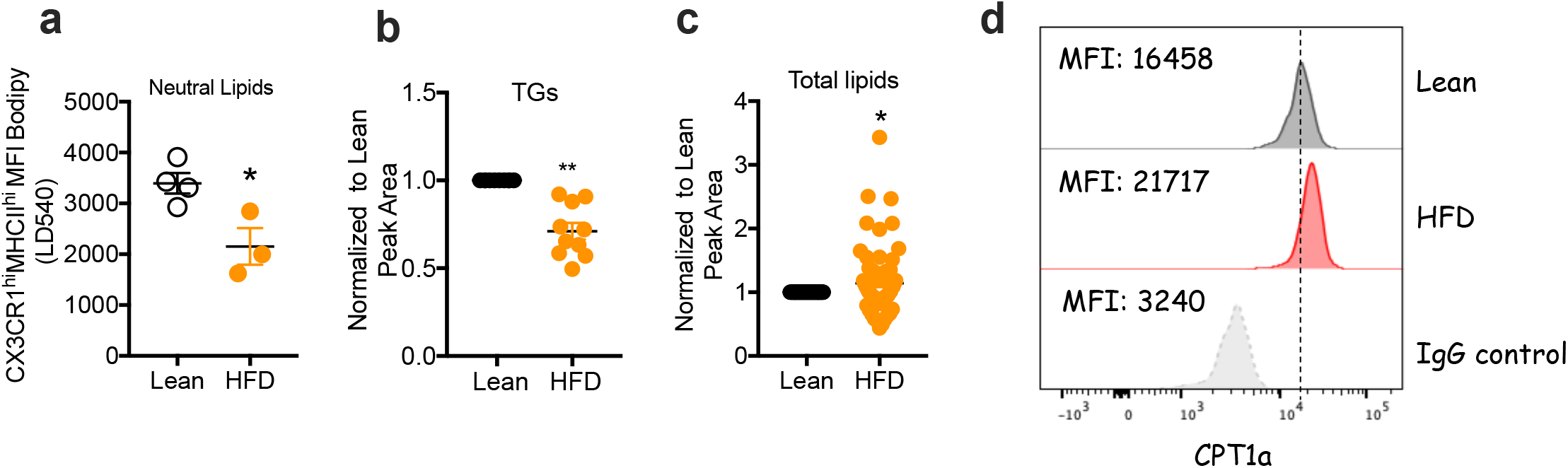
CX3CR1^hi^ MHCII^hi^ LPMs from HFD fed mice show decreased neutral lipids accumulation and increased fatty acid oxidation. **(a)** neutral lipid accumulation in CX3CR1^hi^ MHCII^hi^ LPMs detected using LD540 staining. **(b)**Total triacylglycerol (TG) in 50.000 sorted CX3CR1^hi^ MHCII^hi^ LPMs analyzed by lipidomics. **(c)** Total lipid content in 50.000 sorted CX3CR1^hi^ MHCII^hi^ LPMs analyzed by lipidomics. **(d)** Flow cytometry analysis CPT1a expression in CX3CR1^hi^ MHCII^hi^ LPMs. ∗p < 0.05; ∗∗p < 0.01.

### HFD feeding increases colonic resident macrophage phagocytic capacity

A major recognized function of resident colonic LPMs is phagocytosis related to the maintenance of tissue homeostasis [15]. Indeed, we found that phagocytosis and efferocytosis signatures were enriched in cluster 2 (Figure 4b). We asked whether we could measure this function of LPMs to conclude that it was affected by HFD feeding. We found that colonic CX3CR1^hi^ MHCII^hi^ LPMs from HFD fed mice had increased phagocytic capacity compared to the equivalent cells from lean mice (Figure 6a-b). This was associated with increased reactive oxygen species (ROS) (Figure 6c), which corroborates increased phagocytosis [35]. Overall, our findings indicate that during HFD-feeding a population of CX3CR1^hi^ MHCII^hi^ macrophages develops in the LP that resembles LAMs. This population exhibits enhanced phagocytic capacity.

**Figure 6:**
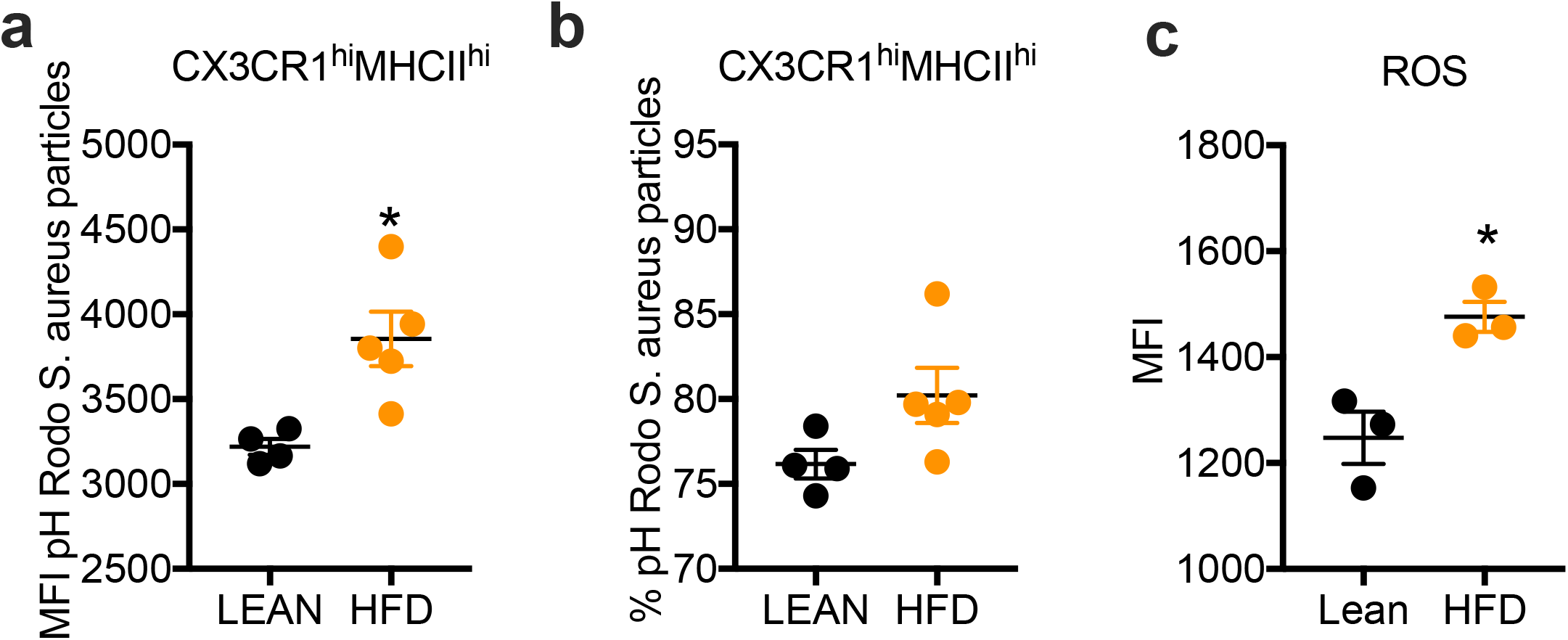
CX3CR1^hi^ MHCII^hi^ LPMs from HFD fed mice have increased phagocytic capacity. **(a-b)** Phagocytosis assay using pH Rodo S. aureus particles in CX3CR1^hi^ MHCII^hi^ LPMs from Lean and HFD fed mice (MFI) and **(b)** percentage. **(c)** Cellular ROS measured ex vivo in CX3CR1^hi^ MHCII^hi^ LPMs from Lean and HFD fed mice. ∗p < 0.05

## Discussion

We have found that HFD increases the percentage and total number of CD64^+^CD11b^+^ macrophages in the intestine, and that a component of this response is comprised of an increase in CX3CR1^hi^ MHCII^hi^ colonic resident LPMs. We speculate that this reflects increased replenishment by blood monocytes which would be appropriate for the intestinal immune system, where constant exposure to bacteria and other materials requires rapid and aggressive responses to potential pathogens to maintain homeostasis, especially in obesity where gut permeability is increased. It is not surprising that we do not see a decrease in the CX3CR1^hi^ MHCII^hi^ population, since the inflammation induced in obesity by HFD is a systemic, chronic low-grade inflammation which does not disrupt the processes that normally induce the full differentiation of resident macrophages [15, 16]. Intestinal resident macrophages have a role in maintenance of tissue homeostasis, inflammation and resolution after inflammation. During intestinal inflammation, the terminal differentiation of monocytes into mature resident macrophages (CX3CR1^hi^) is disrupted [16]. We found that HFD is not sufficient to disrupt differentiation, but rather promoted increased differentiation towards the MHCII^hi^CX3CR1^hi^ phenotype. It is unclear whether this reflects loss of intrinsic factors that usually specify full maturation, or if chronic low-grade inflammation actively revises these processes. We postulate that these might be associated with the chronic low-grade inflammation induced by HFD and increased lipid metabolism, and is an adaptation that prevents inflammatory bowel disease development in this setting.

Previous studies have found that innate gut immunity is involved in metabolic disease as mice fed a HFD have elevated levels of TLR4, TNF, and NF-kB in the gut [36, 37], and increased proinflammatory colonic macrophage infiltration [19]. This agrees with our idea of increased infiltration and differentiation of blood monocytes into resident macrophages. In obese humans, an increase in the number of leukocytes in the intestinal mucosa and a change toward proinflammatory macrophages have been found and these proinflammatory macrophages are potentially recruited via blood monocytes [6, 9, 38]. Our data indicates that the inflammation induced by HFD requires recruitment of blood monocytes to differentiate into CX3CR1^hi^MHCII^hi^ macrophages.

We found that HFD increases protein expression of CD86 and CD206 by CX3CR1^hi^ MHCII^hi^ macrophages, however, no significant changes were observed at mRNA levels. We speculate that the protein levels are more conserved than mRNA levels of these genes. The increased CD86 expression suggests increased co-stimulation of T cells by these resident macrophages [39] and the expression of CD206 indicates that these cells present a mature phenotype and suggests an antiinflammatory phenotype, in agreement with the role of resident macrophages, highly phagocytic without inducing local inflammation [14, 40]. This suggests that upon HFD, CX3CR1^hi^ MHCII^hi^ LPMs maintain their antiinflammatory profile in order to keep intestinal homeostasis. Consistent with this, our scRNAseq shows that HFD results in an increased proportion of a cluster of highly phagocytic related macrophages (Cluster 2). Rohm et al [38] found, in the human intestine, five populations of macrophages: a monocyte derived, a proinflammatory population able to secrete proinflammatory cytokines [16, 18, 41] and two resident/mature (antiinflammatory) macrophages populations. The inflammatory population was increased in all sections of the intestine of obese individuals. Furthermore, increased proinflammatory macrophages in the gut of mice fed a HFD were reported previously [38], and colon-specific macrophage depletion improved glucose tolerance and insulin sensitivity [38]. However, our single cell data is a more robust characterization of heterogeneous macrophage populations in the gut.

scRNAseq allowed us to define a macrophage population with increased phagocytosis/efferocytosis and lipid metabolism. Functionally, LAMs were characterized by a canonical functional signature of lipid metabolism, enhanced phagocytosis and elements of M2 activation [20, 24], which is also observed in our model. LAMs were first described as necessary for controlling metabolic homeostasis [20]. Depletion of these cells caused increased adipocyte hypertrophy, systemic hypercholesterolemia, inflammation and glucose intolerance in mice [20]. Moreover, LAMs were also found to be induced by local lipid exposure in steatotic regions of the murine and human liver [42]. In agreement, in non-alcoholic steatohepatitis (NASH), TIM4^+^ Kupfer cells (KCs) decrease and TIM4^−^ monocyte-derived macrophages are increased in the liver. This monocyte-derived macrophage population consists of cells expressing *Cx3cr1/Ccr2* and cells expressing LAM markers which were associated with crown-like structures and appear to protect the liver against adverse liver remodeling during NASH, preventing liver fibrosis [21].

We also found that under HFD, long-lived resident LPMs that express CD4 and TIM4 are depleted from the colonic LP, as well as decreased gene expression of *Timd4*. These cells correspond to a small fraction of self-maintaining macrophages that are independent of blood monocytes [13, 14]. HFD seems to accelerate the loss of these cells from the intestine, something that normally occurs as mice age [13]. In addition, lipotoxicity would also impact on their survival, since HFD confers a lipid rich environment, especially in adipose tissue macrophages [43], however, we do not observe accumulation of lipid droplets in CX3CR1^hi^ MHCII^hi^ LPMs. Moreover, we found increased numbers of CD4^−^TIM4^−^ macrophages in colonic LP in obese mice. Elsewhere, the CD4^−^TIM4^−^ macrophage population in the colonic LP was reported to be completely dependent on blood monocyte replenishment [13]. The increased resident LPM infiltration upon HFD might be related to an intrinsic mechanism to maintain their survival to support phagocytosis.

The LAM associated signature found in the phagocytic macrophage cluster, which is increased in HFD-fed mice may be necessary for homeostasis of the gut during obesity. This seems to be associated with balancing lipid accumulation. Consistently, we found decreased lipid droplet accumulation in CX3CR1^hi^MHCII^hi^ LPMs from HFD-fed mice. This reflected decreased TG accumulation and increased expression of genes related to fatty acid oxidation and lipolysis, which was also associated with increased CPT1a protein expression by these cells, supporting the idea that there is increased fat catabolism in these cells. Inflammatory macrophages were shown to accumulate LD because they are not able to oxidize lipids and this accumulation is necessary for proinflammatory cytokine release [44]. However, efficient phagocytosis was also shown to require fatty acids hydrolysis [45] and LPMs are known to be highly phagocytic [14], patrolling the gut to maintain homeostasis. Moreover, we detected increased ROS in CX3CR1^hi^MHCII^hi^ LPMs, consistent with increased phagocytosis. Corroborating, we found an increased efferocytosis signature in the macrophage cluster that was increased in HFD lamina propria. It was previously shown that efferocytosis of apoptotic cells by macrophages anchor the resolution of intestinal inflammation, preventing necrosis and further inflammation, and programing macrophages for tissue repair [46, 47]. Moreover, efferocytosis by intestinal macrophages is dependent on COX2 [46], which is necessary for prostaglandins synthesis [48]. Prostaglandin synthesis is necessary for phagocytosis in inflammatory macrophages [44], supporting the role of lipid metabolism and LAMs in contributing to homeostasis in the gut during HFD. In addition, we postulate that the increased phagocytosis might reflect the increased prevalence of cluster 2 macrophages in mice fed HFD.

Our study provides a new perspective on the phenotype of resident LPMs upon HFD feeding and obesity. We conclude that HFD induces changes in the populations profile of colonic resident LPMs, increasing activation, depleting long lived CD4^+^TIM4^+^ LPMs, and increasing monocyte-derived LAMs displaying of increased phagocytosis capacity in order to maintain gut homeostasis during HFD feeding.

## Methods

### High fat diet treatment

Obesity was induced by ad libitum feeding of C57/BL6 CX3CR1 GFP heterozygous mice for 12 weeks with irradiated high fat diet (Rodent Diet 60% kcal from fat, Research Diets, Inc., Cat #D12492). Control diet (chow) containing 24% protein, 47.5% carbohydrate, and 4.9% fat, was given to age and sex matched animals as a control group.

### Metabolic Parameters Analysis

Peripheral response to glucose was assessed by glucose tolerance test (GTT). Glucose (2g/Kg body weight) was administered intraperitoneally in mice fasted for 12 hours. Glucose levels in blood were determined before and after 15, 30, 60, 90 and 120 minutes from glucose administration. The insulin response (insulin tolerance test (ITT) was examined after fasting mice for 6 hours. Blood glucose levels were determined before and after 15, 30, 60, 90 and 120 minutes of 0.8 U/kg insulin administration. All mice were maintained in individual cages. It was used ACCU-CHEK®Advantage II - (Roche Mannheim, Germany) for reading the blood glucose levels.

### Determination of Serum LPS

LPS concentration in serum was determined using a chromogenic assay based on a Limulus amebocyte extract (LAL kit-terminal QCL1000) (LONZA, Basel, Switzerland). Samples were collected at the time of euthanasia via cardiac puncture to try to minimize as much as possible the chance of contamination. The test is quantitative for Gram-negative bacterial endotoxin and was carried out according to manufacturer’s guidelines. We used 50 microliters of samples in duplicates. The kit’s sensitivity is 0.1 endotoxin units/mL.

### Lamina propria cells/macrophages isolation

At the end of treatment, animals were sacrificed, and cells from large intestine lamina propria recovered. Briefly, the large intestine was separated at the junction with the cecum, and all remaining connective and fat tissue removed. Isolated intestine was then opened longitudinally, cleaned, cut into 0.4-1 cm pieces and washed with ice-cold 25 mM HEPES i n PBS. Tissue fragments were then placed in RPMI (Gibco) medium containing 3% fetal bovine serum, 25 mM HEPES, 5 mM EDTA plus 3.5 mM Dithiothreitol and incubated for 15 min in 37°C with gentle agitation. Tissue fragments were recovered via filtering and vigorously washed thrice with 2 mM EDTA i n RPMI, discarding supernatants. Washed fragments were minced then incubated i n RPMI supplemented with 0.5% fetal bovine serum, 2 mg/mL Collagenase VIII (Roche) and 0.5 mg/mL DNAse I (Roche) for 30-40 min at 37ºC with gentle agitation. Cell suspension and tissue fragments were filtered through a 70 μm strainer, and dissociated with the rubber end of a syringe plunger. Resulting cell suspension was centrifuged and further filtered through a 40 μm cell strainer. Finally, live cells were passed through a Percoll gradient (35%/70%), recovered from the interface, washed and counted.

### RNA sequencing

Total RNA was extracted with the RNAqueous-Micro Total RNA Isolation kit (Thermo Fisher Scientific) and quantified using Qubit 2.0 (Thermo Fisher Scientific), according to the manufacturer’s instructions. Libraries were prepared using the TruSeq stranded mRNA kit (Illumina) and sequenced in a HISeq 3000 (Illumina) by the Deep Sequencing Facility at the Max Planck Institute for Immunobiology and Epigenetics. Sequenced libraries were processed with a pipeline optimized by the Bioinformatics core at the Max Planck Institute for Immunobiology and Epigenetics [49]. Raw mapped reads were processed in R (Lucent Technologies) with DESeq2 [50] to determine differentially expressed genes and generate normalized read counts to visualize as heatmaps using Morpheus (Broad Institute). Enrichment analysis based on hypergeometric distribution followed by false discovery rate (FDR) correction for GO terms were made using ShinyGO 0.76 software [51].

### Single cell barcoding and library preparation

Lamina propria cells were prepared for cell sorting as described above, using only LIVE/DEAD dye and an anti-CD45 (BioLegend, clone: 30-F11) fluorochrome-conjugated antibody. Recovered CD45 positive cells were prepared for scRNAseq analysis using a 10X Genomics Chromium Controller. Single cells were processed with GemCode Single Cell Platform using GemCode Gel Beads, Chip and Library Kits (v2) following the manufacturer’s protocol. Libraries were sequenced on HiSeq 3000 (Illumina). Samples were demultiplexed and aligned using Cell Ranger 2.2 (10X Genomics) to genome build GRCh38 to obtain a raw read count matrix of barcodes corresponding to cells and features corresponding to detected genes. Read count matrices were processed, analyzed and visualized in R using Seurat v. 3 and Uniform Manifold Approximation and Projection (UMAP) as a dimensionality reduction approach. Differentially expressed genes, with greater than a 1.2 fold change and an adjusted p value of less than 0.05, were obtained and compared across clusters using the UpSet methodology. Unbiased cell classification was based on scGate [32] annotation to evaluate the strength of signature marker expression in each cell. Gene expression scores were calculated using UCell. The LAM signature was based on Jaitin et al, 2019 [20]. The efferocytosis signature was based on a GO term. OXPHOS and Glycolysis are based on KEGG. Heat shock and Ifn signatures were based on an r Package from the Carmona lab in Lausanne (UCell score creators).

### Lipidomics

The protocol for lipid extraction was adapted from Matyash et al [52]. Frozen cell pellets (5x 10^5^ cells) were resuspended in ice cold PBS and transferred to glass tubes before the addition of methanol and methyl tert-butyl ether. The tubes were then shaken for 1 h at 4 °C. Water was added to separate the phases before centrifugation at 1000 g for 10 min. The upper organic phase was collected and dried in a Genevac EZ2 speed vac. Samples were resuspended in 2:1:1 isopropanol: acetonitrile:water prior to analysis. LC–MS was carried out using an Agilent Zorbax Eclipse Plus C18 column using an Agilent 1290 Infinity II UHPLC in line with an Agilent 6495 Triple Quad QQQ-MS. Lipids were identified by fragmentation and retention time, and comparison to standards, and were quantified using Agilent Mass Hunter software. Comparisons were made between relative amounts of lipid between conditions, extracted from equivalent cell numbers. Peak areas were quantile-normalized across the batch to generate the lipid intensities used for the plots and subsequent statistics shown in this manuscript.

### Flow cytometry

After lamina propria cells isolation from CX3CR1 GFP heterozygous mice, the cell suspension was incubated in 5 μgml−1 anti-CD16/32 (#101302, Biolegend, 1:200), stained with Live Dead Near IR (#L10119,ThermoFisher, 1:500), and then surface stained with a fluorochrome conjugated monoclonal antibody to CD45 (Pacific Blue, Biolegend, clone SF18009F, 1:200), CD11b (PECy7, Biolegend, clone M1/70,1:300), CD64 (BV711, Biolegend, X54-5/7.1, 1:200), MHCII (PercpCy5.5, Biolegend, M5/114.15.2, 1:200), Tim4 (APC, Biolegend, clone: F31-5G3, 1:200), CD4 (BV510, Biolegend, clone: RM4-5, 1:200), CD86 (BV650, Biolegend, clone: GL-1, 1:400), CD80 (APC, Biolegend, clone 16-10A1, 1:200), CD206 (PE, Biolegend, clone C068C2, 1:500). For Phagocytosis assay, pHrodo™ Red Staphylococcus aureus Bioparticles™ (#A10010, ThermoFisher, 100μg ml−1) was used according to the manufacturer’s instructions. BODIPY LD 540 (developed by Prof. Christoph Tiele [53] was used for neutral lipid staining according to the manufacturer’s instructions. CPT1a primary antibody staining (Proteintech Group Cat# 15184-1-AP RRID: AB_2084676) was detected using the Alexa Fluor-647 anti-rabbit secondary antibody (Life technologies). ROS was measured using CellROX Deep Red Flow cytometry assay Kit according to manufacturer instructions (Invitrogen #C10491). Data were acquired by flow cytometry on an LSRII or LSR Fortessa (BD Biosciences) and analyzed with FlowJo v.10.1 (Tree Star).

### Statistical analysis

Statistical analysis was performed using prism 7 software (Graph pad) and results are represented as mean ± s.e.m. Comparisons for two groups were calculated using unpaired two-tailed Student’s t tests, comparisons of more than two groups were calculated using one-way ANOVA with Bonferroni’s multiple comparison tests. We observed normal distribution and no difference in variance between groups in individual comparisons. Statistical significance: ∗p <0.05; ∗∗p < 0.01; ∗∗∗p < 0.001; ∗∗∗∗p < 0.0001. Further details on statistical analysis are listed in the figure legends.

### Data availability

Single Cell RNA-sequence data have been deposited in Gene Expression Omnibus under the primary accession code GSE171330.

## Acknowledgments

We thank the members of the Pearce laboratories, the FACS, and Deep Sequencing facilities at the Max Planck Institute of Immunobiology and Epigenetics, the members of Camara’s Laboratory, and CEFAP-USP for technical support. A.C. was supported by the CAPES/Alexander von Humboldt Fellowship Foundation (88881.136065/2017-01) and FAPESP grants 2015/18121-4, 2017/05264-7 and 2017/00721-0. This study was also financed in part by CAPES, finance code 001.

## Author Contributions

A.C, E.J.P. and N.O.S.C. conceptualized the study, designed the research, and interpreted the data. D.E.S, T.A.S. and N.R analyzed bioinformatics data. A.C, J.E-H., J.B. analyzed Lipidomics data. A.C, N.v.T.B., C.F.A., L.B.M., N.R., K.M.G, A.M.K, J.C., A.M.C., G.C., T.A.S., F.O.S. performed experiments. E.L.P. helped with project insights. N.O.S.C., E.L.P. and E.J.P. acquired funding. A.C., E.J.P. and NOSC wrote the manuscript.

## Figure Legends

**Supplemental Figure 1:**
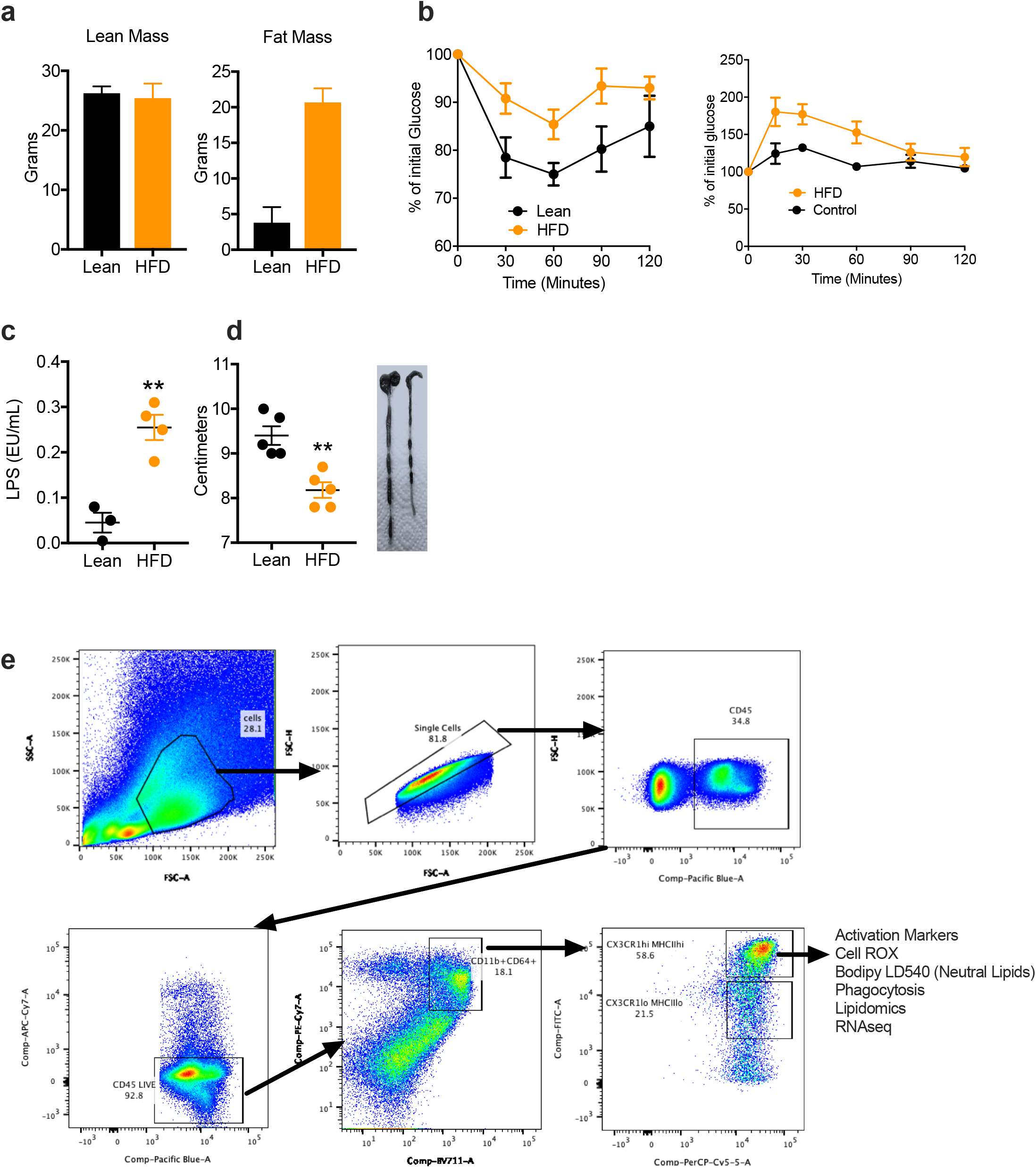
Metabolic parameters after 12 weeks of chow (Lean) or HFD. **(a)** Lean mass and Fat mass in Lean and HFD fed mice measured by Eco-MRI after 12W of Chow or HFD. **(b)** Insulin resistance test and glucose tolerance test performed 12 weeks after chow (Lean) and HFD. **(c)** Serum LPS from Lean and HFD fed mice. **(d)** Size (cm) of the colon measured after 12 weeks after chow (Lean) and HFD. **(e)** Gating strategy used to identify CX3CR^hi^ MHCII^hi^ colonic LPMs. ∗∗p < 0.01.

**Supplemental Figure 2:**
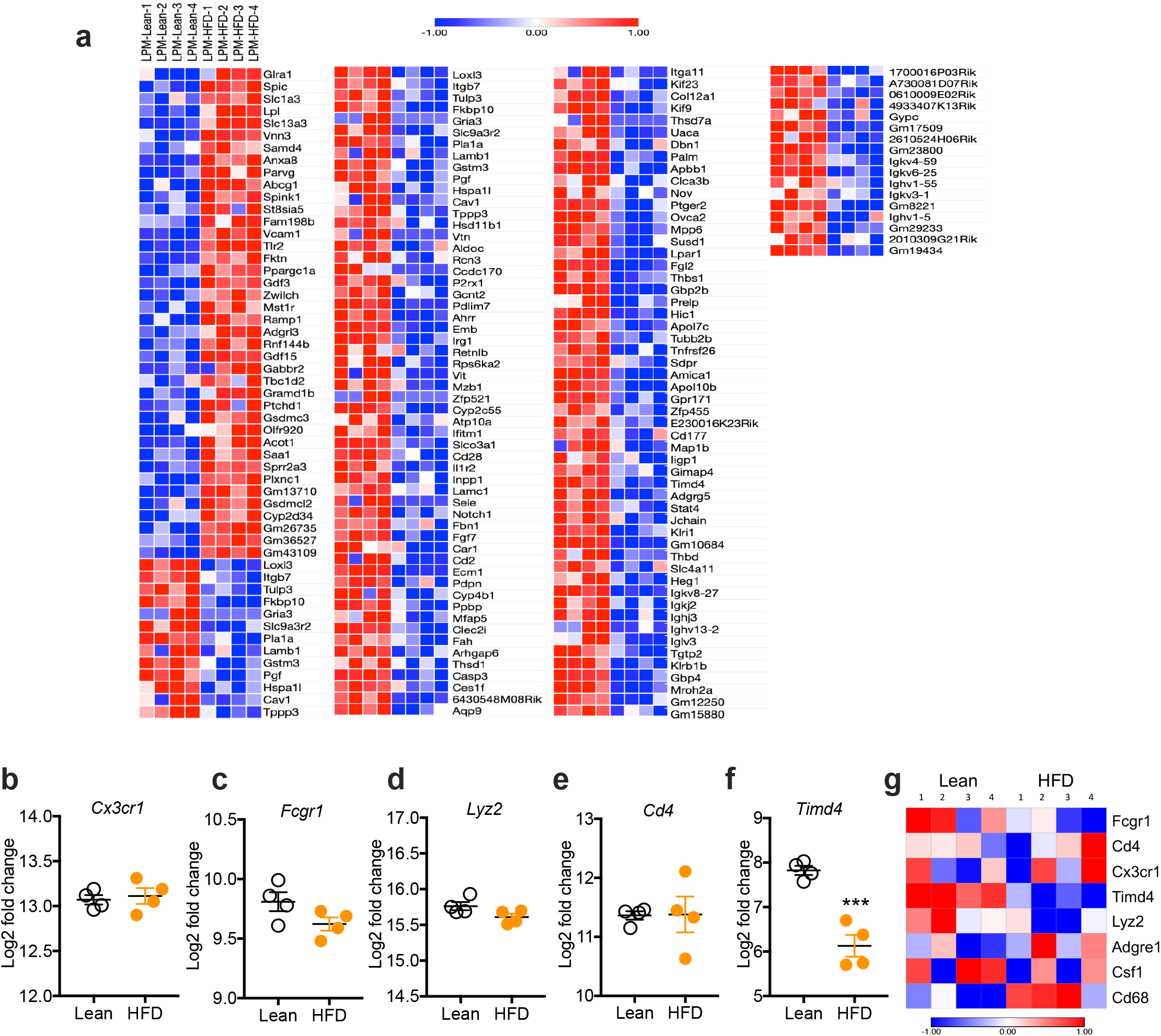
12 weeks of chow (Lean) or HFD modulate gene expression in colonic LPMs. **(a)** 167 regulated genes (>1 Fold Change) in sorted CX3CR1^hi^ MHCII^hi^ LPMS. **(b-g)** Log2 Fold change of gene expression of *Cx3cr1, Fcgr1, Lyz2, Cd4 and Timd4* in sorted CX3CR^hi^ MHCII^hi^ LPMs by RNA sequencing. ∗∗∗p < 0.001.

**Supplemental Figure 3:**
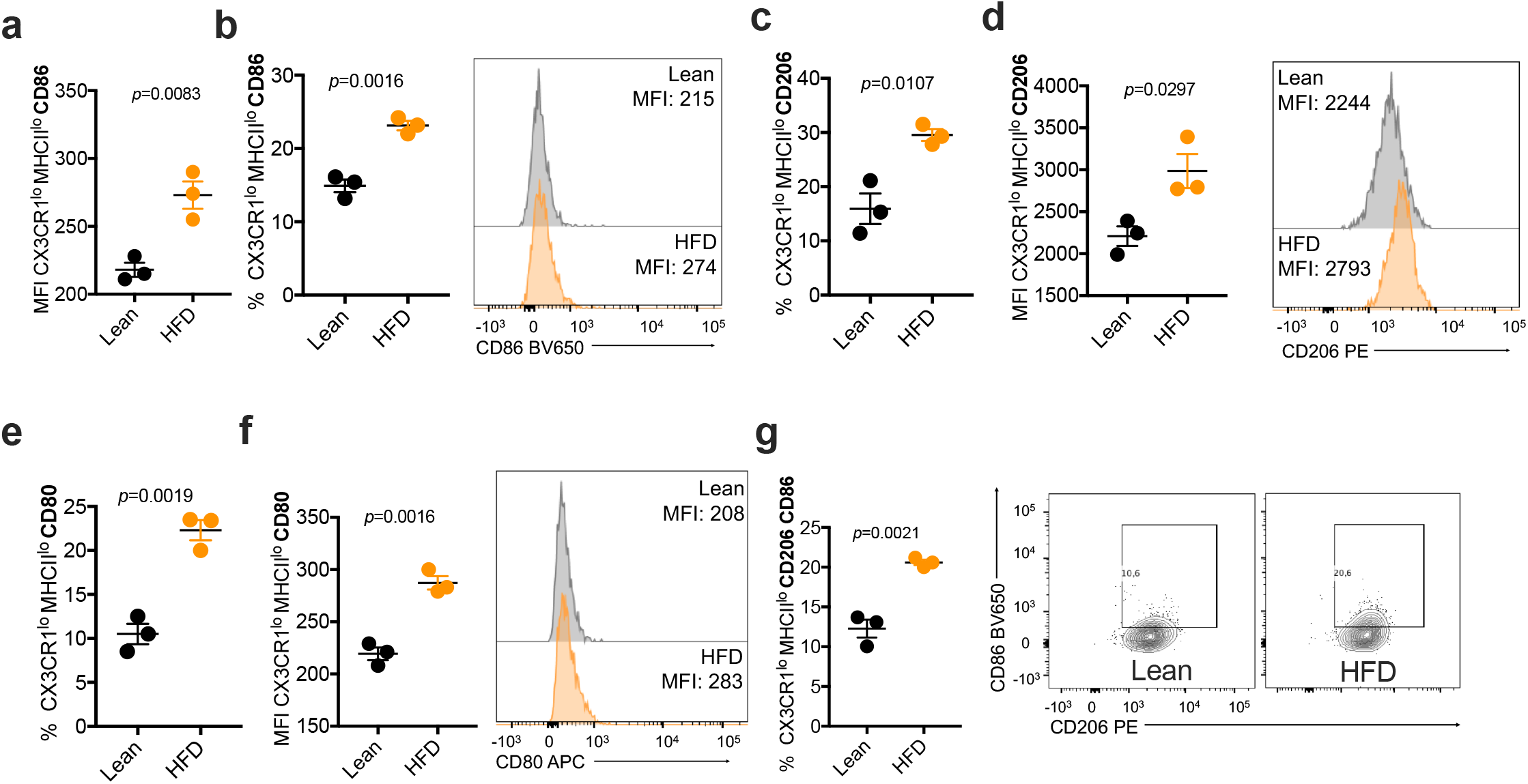
HFD modulates CD86, CD80 and CD206 in MHCII ^lo^CX3CR^lo^ LPMs. **(a)** Percentage of CX3CR1^lo^ MHCII^lo^ LPMs expressing CD86 and **(b)** MFI of CD86. **(c)**Percentage of CX3CR1^lo^ MHCII^lo^ LPMs expressing CD80 and **(d)** MFI of CD80. **(e)**Percentage of CX3CR1^lo^ MHCII^lo^ LPMs expressing CD206 and **(f)** MFI of CD206. **(g)**Percentage of CX3CR1^lo^ MHCII^lo^ LPMs expressing CD206 CD86.

**Supplemental Figure 4:**
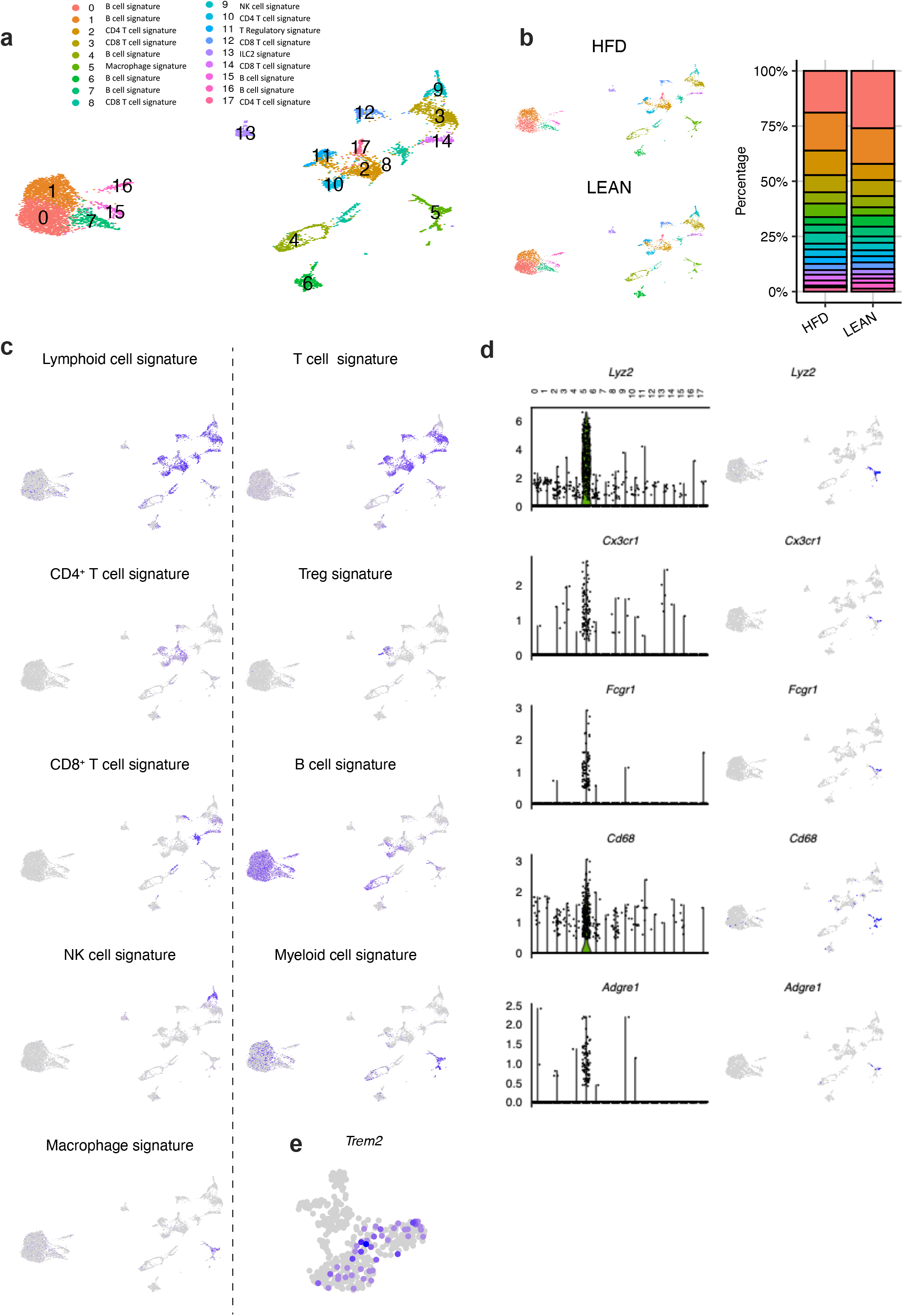
Single cell RNA sequencing on CD45+ cells isolated from colonic LP. **(a-b)** Single cell RNA sequencing of CD45+ cells from the LP isolated from 6 mice (3 (12W) HFD and 3 lean) and loaded onto the 10X Genomics Chromium platform. 17 clusters were identified. **(c)** Unbiased cell classification, based on scGate. (d) Violin plots showing *Lyz2, Cx3cr1, Fcgr1, Cd68* and *Adgre1* increased expression in cluster 5 and scatter plots of macrophage genes. **(e)** Scatter plot showing Trem2 expression by clusters 1, 2 and 3.

**Supplemental Figure 5:**
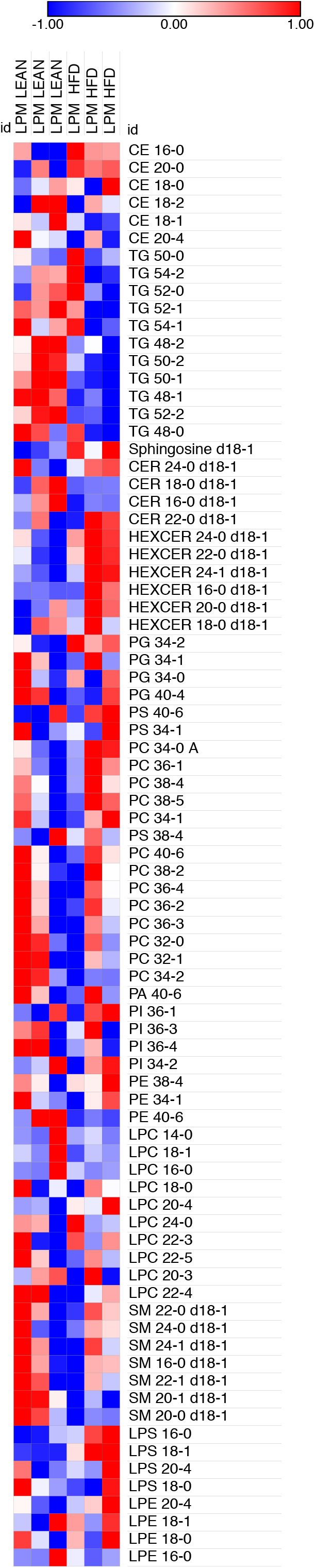
Lipidomics analysis on CX3CR1^hi^ MHCII^hi^ LPMs from Lean and HFD-fed mice. Lipid species identified on CX3CR1^hi^MHCII^hi^ LPMs isolated from colon of Lean and HFD-fed mice.

